# SARS-CoV-2 entry sites are present in all structural elements of the human glossopharyngeal and vagal nerves: clinical implications

**DOI:** 10.1101/2021.12.30.474580

**Authors:** L. Vitale-Cross, I Szalayova, A Scoggins, M. Palkovits, E Mezey

## Abstract

Severe acute respiratory syndrome coronavirus (SARS-CoV-2) infections result in the temporary loss of smell and taste (anosmia and dysgeusia) in about one third of confirmed cases. Several investigators have reported that the viral spike protein receptor is present in olfactory neurons. However, no study has been published to date showing the presence of viral entry sites angiotensin-converting enzyme 2 (ACE2), neuropilin1 (NRP1), and TMPRSS2, the serine protease necessary for priming the viral proteins, in human nerves that are responsible for taste sensation (cranial nerves: VII, IX and X). We used immunocytochemistry to examine three postmortem donor samples of the IXth (glossopharyngeal) and Xth (vagal) cranial nerves where they leave/join the medulla from three donors to confirm the presence of ACE2, NRP1 and TMPRSS2. Two samples were paraffin embedded; one was a frozen sample. In addition to staining sections from the latter, we isolated RNA from it, made cDNA, and performed PCR to confirm the presence of the mRNAs that encode the proteins visualized. All three of the proteins required for SARS-CoV-2 infections appear to be present in the human IXth and Xth nerves near the medulla. Direct infection of these nerves by the COVID-19 virus is likely to cause the loss of taste experienced by many patients. In addition, potential viral spread through these nerves into the adjacent brainstem respiratory centers might also aggravate the respiratory problems patients are experiencing.

## Introduction

Since the start of the Covid19 virus pandemic two years ago in 2019 more than 250 million people have been infected and over 5 million have died worldwide. Information about the virus has grown at an amazingly fast pace. The expectation that the number of infected people might be 50–80% of the world’s population suggests that the overall number of patients with neurological disease could become rather significant [1]. The widely distributed ACE2 receptor was identified as the primary binding site [2] for the virus, and later TMPRSS2, an intracellular protease, was shown to promote viral entry into cells by cleaving the S protein into S1 (receptor binding) and S2 (membrane fusion domains). The latter mediates host-virus fusion. The membrane protein neuropilin1 (NRP1) is an alternate entry site for the virus [3]. Data regarding entry proteins led to the development of antiviral vaccines and therapeutic agents. In parallel with these efforts, we have learned a lot about the symptoms of COVID-19 and how to differentiate it from other viral diseases. Among the unique symptoms of the infection are loss of taste and smell in about one-third of patients and papers have been published describing the cause of these symptoms. It has been shown that molecules and pathogens can migrate across the cribriform plate (paracellular migration) [4,5] from the infected olfactory epithelium. The perineural channels yield a direct connection to the cerebrospinal fluid (CSF) space and the olfactory bulb. These channels have been known for centuries to connect the nasal cavity to the central nervous system (CNS) extracellular/CSF space see [6]. Even though we may understand how SARS-CoV-2 makes its way to the CSF space [7], we lack data about the exact location of viral entry sites responsible for taste-loss in Covid-19 infected patients. Thus, we set out to use samples of the glossopharyngeal and vagal cranial nerves from postmortem human samples to see which cellular structures of these nerves’ express receptors for the Covid-19 SARS virus.

## Methods

### Brain and Nerve Samples

Anonymized postmortem human brain samples were obtained from the Human Brain Tissue Bank, Semmelweis University, Budapest, Hungary. The brains were dissected and specific brainstem areas with cranial nerves were isolated. The samples were either flash frozen or embedded in paraffin after formalin fixation. The study conformed to European ethics regulations (TUKEB #189/2015).

### Immunocytochemistry (ICC)

The paraffin-embedded sections were deparaffinized with SafeClear II (Fisher Scientific; #044-192) and rehydrated in decreasing concentrations of ethanol followed by heat-induced epitope retrieval (HIER) in 10 mM citrate buffer at pH 6.0 in a microwave oven. Slides were placed in the oven lying flat in a plastic container and covered with the citrate buffer. They were brought to boil at high power (700 W), and then incubated for 5 more minutes at 50% power (350 W). After HIER, the slides were allowed to cool to room temperature in the buffer. Next, Bloxall (Vector; SP-6000) dual endogenous enzyme blocking solution was applied to the sections for 15 min before the primary antibodies were added. Primary antibodies used: ACE2 (rabbit), Abcam ab15348; NRP1 pre-conjugated to Alexa-488; Neurofilament (chicken), Abcam ab4680; TMPSSR2 (rabbit), Novus Biologicals NBP3-00492; MBP pre-conjugated to Alexa-647; fibronectin (rabbit) gift from Dr. Ken Yamada ([8]. Secondary antibodies were purchased from Jackson Laboratories, raised in donkey, anti-rabbit (Cat#711-586-152) or anti-chicken (Cat#ab150170). For more details about the stainings see Fig. legends. When we needed to stain with multiple antibodies from the same species, we used a multiplex labeling method based on signal amplification and fluorescent tyramide dyes [9]. The advantage of this technique is that antibodies from the same species can be used consecutively because the tyramide-conjugated fluorescent dye is insoluble in water allowing both the primary and secondary antibodies to be removed by heat. The fluorescent signal from each insoluble tyramide complex remains where the target antigen was. This process can be replicated several times using different fluorochromes conjugated to tyramide. The fluorescent signal emitted by the HRP–tyramide complex is much stronger than the one a traditional fluorochrome-conjugated secondary antibody gives. Briefly, the slides were incubated overnight at 4 °C with the first primary antibody followed by an anti-IgG for the appropriate species. The IgGs were preconjugated to an HRP polymer (VisUCyte HRP polymer; R&D Systems). The signal was then visualized by adding different color fluorochrome tyramide conjugates, which are high-affinity substrates of the HRP. After staining with the first primary antibody, the microwave cycle was repeated, leaving only one specific tyramide signal. Then additional primary antibodies (and fluorochrome tyramide conjugates) were used one after another to visualize the target proteins. For control staining, the primary antibody was omitted, but the amplification process including the incubation with the HRP-polymers and the tyramide-fluorochromes was unchanged. All antibodies were also used for individual unamplified stainings to ensure specificity. After completion of the procedure, all sections were analyzed with a Leica DMI6000 inverted fluorescent microscope using LAX software.

### PCR of human tissue samples

Preparation of cDNA and PCR analysis was performed as follows: Before it was fixed (to perform ICC) a small piece of the frozen sample of the IXth and Xth cranial nerves was broken off the block and placed in Trizol (Thermo Fisher, Waltham, MA). RNA was prepared according to the manufacturer’s protocol. After quantitation by Nanodrop 1.0 μg of RNA was used to prepare cDNA using a BioLink™ cDNA Synthesis Kit, (Washington, DC, catalog number 16-2100-C250). Lung RNA purchased from Ambion (Thermo Fisher, Waltham, MA) was used as a positive control. The cDNA was diluted to 25-50 ng/μl for downstream applications. PCR analysis of the ACE2R, NRP1, TMPRSS2, and ACTB (housekeeping) cDNAs were performed using the Bio-Rad T100 and REDtaq polymerase, (Sigma-Aldrich, St. Louis, MO). The primers used are listed in the table below. The PCR products were run on a 2% agarose gel along with a 100 bp ladder (Thermo Fisher, Waltham, MA).

### Tissue culture

Two immortalized cell lines were used to validate the ACE2 receptor, TMPRSS2 and NRP1 assays: Human oligodendroglioma cells (HOG, Cat# SCC163) were obtained from Millipore Sigma-Aldrich (Billerica, MA) and cultured in DMEM-high glucose medium (Sigma D5796, St. Louis, MO) with 10% fetal bovine serum and antibiotics. Human primary brain vascular fibroblasts (Cat# H-6076) from Cell Biologics, Inc (Chicago, IL). These cells were cultured in complete fibroblast medium (Cell biologics Inc, Cat# M2267) that included all required supplements. For immunostainings the cells were put in chamber slides at 35,000 cells/well and fixed with 4% paraformaldehyde. ICC was performed as above and using two step simple fluorescent immunostaining without microwaving and tyramide amplification. Controls were run with secondary antibodies only.

## Results

### Human nerves

Immunostainings of samples from all three donors showed a very similar picture between the two paraffin embedded and the frozen samples of the IX/X cranial nerves. Also, all sections of the cranial nerve samples examined had similar staining patterns; the branches of both nerves had the same distribution of antigens.

We first used antibodies to identify the two known entry sites for the virus SARS-Cov-2: ACE2 and NRP1, a widely expressed transmembrane glycoprotein that was found to be a co-receptor, facilitating the entry of the virus [10,11]. In addition, we looked for TMPRSS2, a protease shown to “prime” the spike protein to catalyze membrane fusion between the virus and the host cell [12,13]. We used antibodies to stain cellular elements within the nerve bundles: neurofilament protein (NF) to label axons; myelin basic protein (MBP) and myelin protein zero (MP0) to label the myelin; and fibronectin (FN) to label fibrocytes and fibroblasts in and around the nerves. This allowed us to identify which cell types in the nerve express the SARS-Cov-2 entry sites and the protease.

ACE2, the primary viral binding protein, was present in all structures of the nerves, such as nerve sheaths, connective tissue inside the bundles, small vessels, axons and myelin. We then looked at the overlap of ACE2 and NRP1 by staining cross and longitudinal sections of the nerves and found that both labeled connective tissue and the nerve fibers within nerve bundles (Fig.1A,B). ACE2 was in the endoneurium around the myelinated fibers; fine dots of ACE2 staining could be seen in the axon and parts of the myelin sheath as shown by myelin basic protein (MBP) staining. The NRP1 antibody also stained myelin, endoneurium, and connective tissue (Fig. 1B). Vascular wall cross sections were also stained. The proteinase TMPRSS2 was intensely stained in sections of axons and myelin and was less prominent in connective tissue (Fig. 1C). Differences in staining among the different target proteins were observed in different cellular compartments versus different cell types (Fig.1 D). The highest density of ACE2 was in the connective tissue within and surrounding the nerve (Fig. 2A,B, C-inset). Fibronectin (FN) staining depicting connective tissue showed a web-like distribution in a significant overlap with ACE2 (Fig. 1C). TMPRSS2 was present in all cellular elements, but more intense staining was observed in the neural elements compared to the connective tissue and vasculature (Fig. 2D,E) and showed a complementary inverse staining pattern with FN (Fig.2 E-F). The second entry site, neuropilin1 (NRP1) seemed to be more abundant in the neural tissue and in a few connective tissue areas (Fig.2 G-I.).

**Fig.1.**
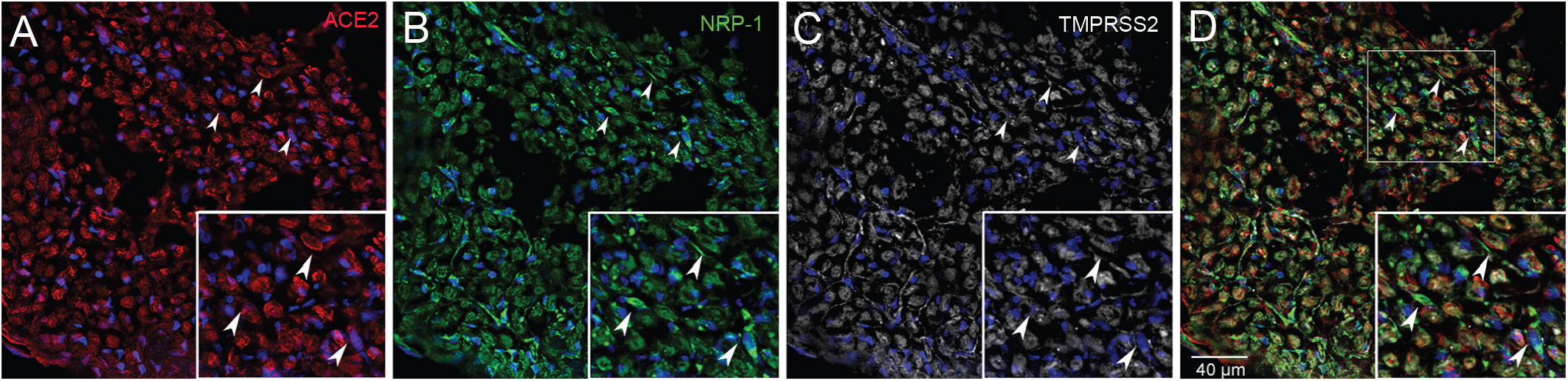
Human glossopharyngeal/vagal nerve at the level of medulla oblongata. Six-micron thin sections were stained for ACE2 (A) using a rabbit primary antibody followed by an Alexa-594 (red) conjugated secondary antibody (Diluted 1:350). Next a mouse primary antibody to TMPRSS2 (C) was applied (Diluted 1:100) followed by an anti-mouse Alexa-647 (colored white) secondary antibody and finally an anti-Neuropilin (B) pre-conjugated to Alexa-488 was used (Diluted 1:100). (D) shows the overlay of the three antibodies. Cell nuclei are stained with DAPI (blue). The inset shows an area enlarged for better visibility. *All primary antibodies were applied overnight.

**Fig.2.**
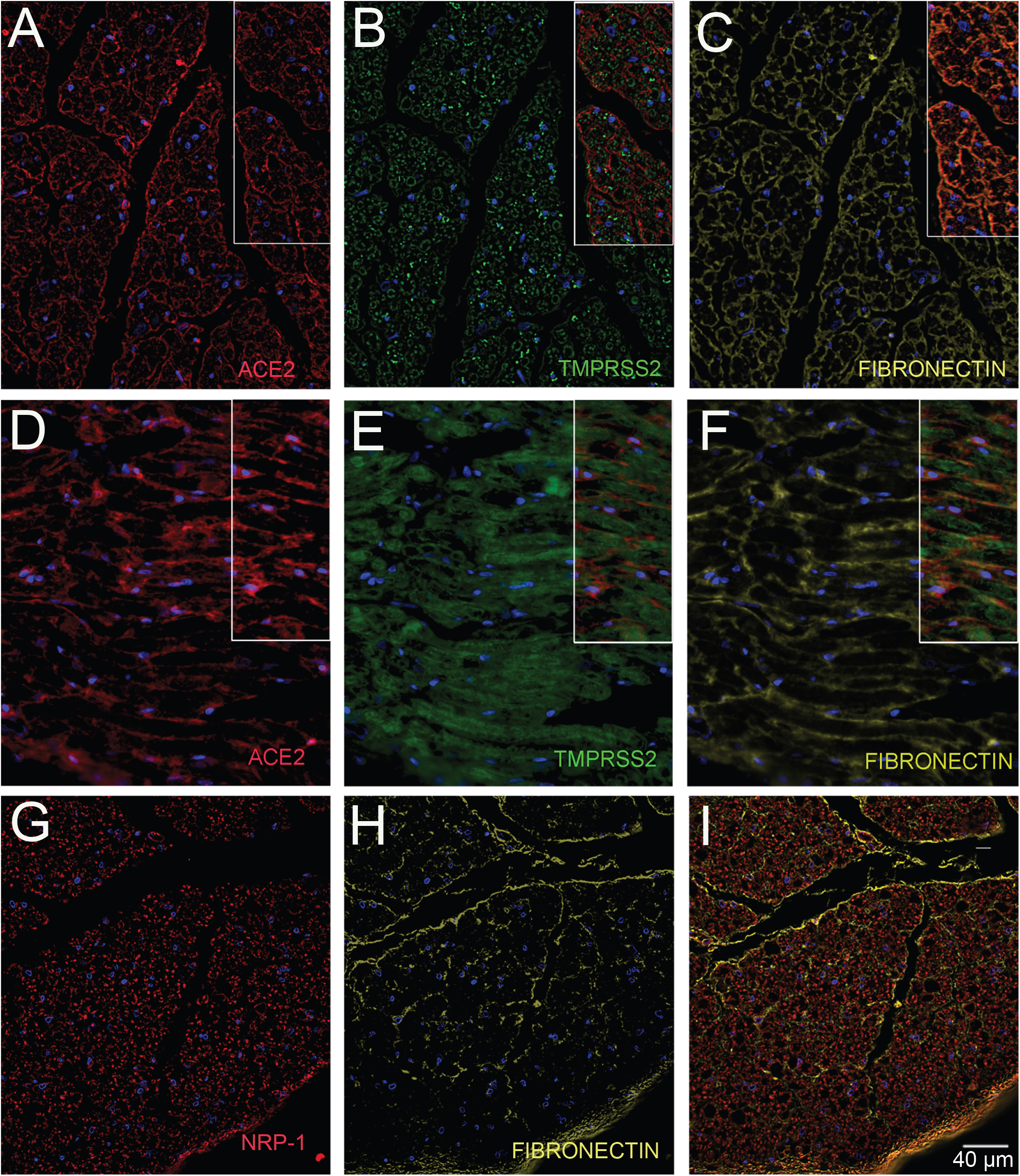
Human glossopharyngeal/vagal nerve at the level of medulla oblongata. The tissue contained both cross sectional cuts of the nerve (A-C & G-I) and longitudinally cut sections of the nerve (D-F). Six microns thin sections were stained for ACE2 (A & D) using a rabbit primary antibody (Diluted 1:350) and amplifying it with Tyramide 594 (Red). Next a rabbit primary antibody to TMPRSS2 (B & E) was applied (Diluted 1:100) followed by a Tyramide 488 amplification (green). At the end, a rabbit primary antibody to Fibronectin (C & F) was applied (Diluted 1:5000) followed up with an anti-mouse Alexa-647 (colored yellow) secondary antibody. A separate section was stained with a rabbit primary antibody to Fibronectin (H) was applied followed by an anti-mouse Alexa-647 (colored yellow) secondary antibody and finally an anti-Neuropilin (G) pre-conjugated to Alexa-488 was used (Diluted 1:100). (I) shows the overlay of the two antibodies. Cell nuclei are stained with DAPI (blue). *All primary antibodies were applied overnight. *All secondary antibodies were diluted 1:500.

Finally, we colocalized ACE2 (Fig. 3A-H) and NRP1 (Fig. 3I-P) in axons, neurofilament (NF) and myelin (MBP) in both longitudinal and cross sections. Both of the above neural elements were positive for both ACE2 and NRP1, suggesting that myelin making cells (oligodendrocytes or Scwann cells) as well as the neurons are both vulnerable for the viral attack by SARS-CoV-2. We summarized our findings and the relative intensity of the staining in Table 1.

**TABLE 1.**
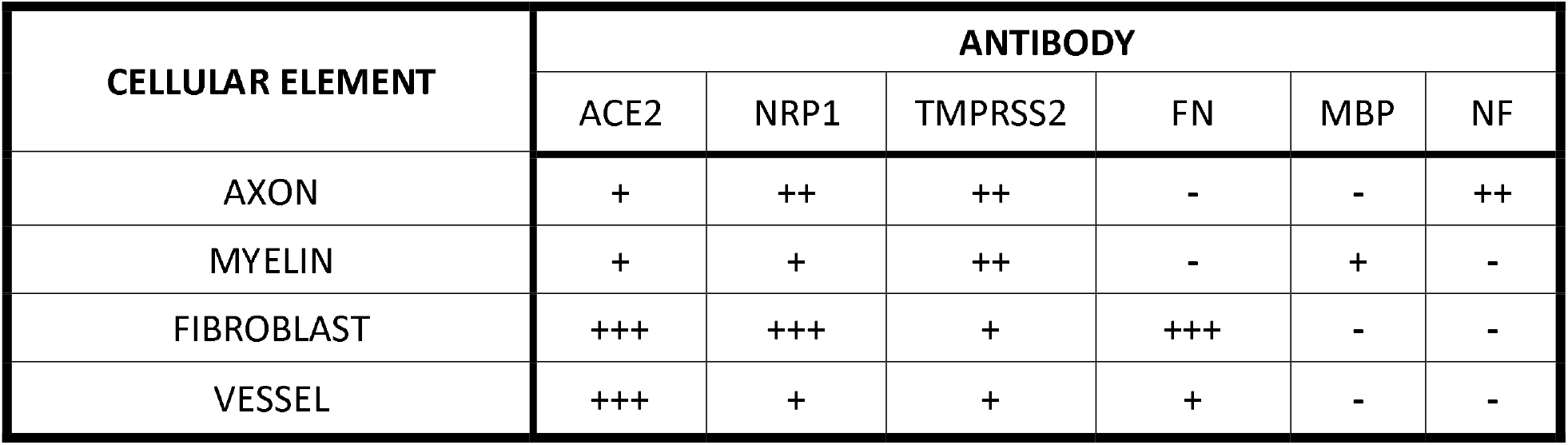
Staining intensity in different cell types

| CELLULAR ELEMENT | ANTIBODY |  |  |  |  |  |
| --- | --- | --- | --- | --- | --- | --- |
|  | ACE2 | NRP1 | TMPRSS2 | FN | MBP | NF |
| AXON | + | ++ | ++ | - | - | ++ |
| MYELIN | + | + | ++ | - | + | - |
| FIBROBLAST | +++ | +++ | + | +++ | - | - |
| VESSEL | +++ | + | + | + | - | - |

**Fig.3.**
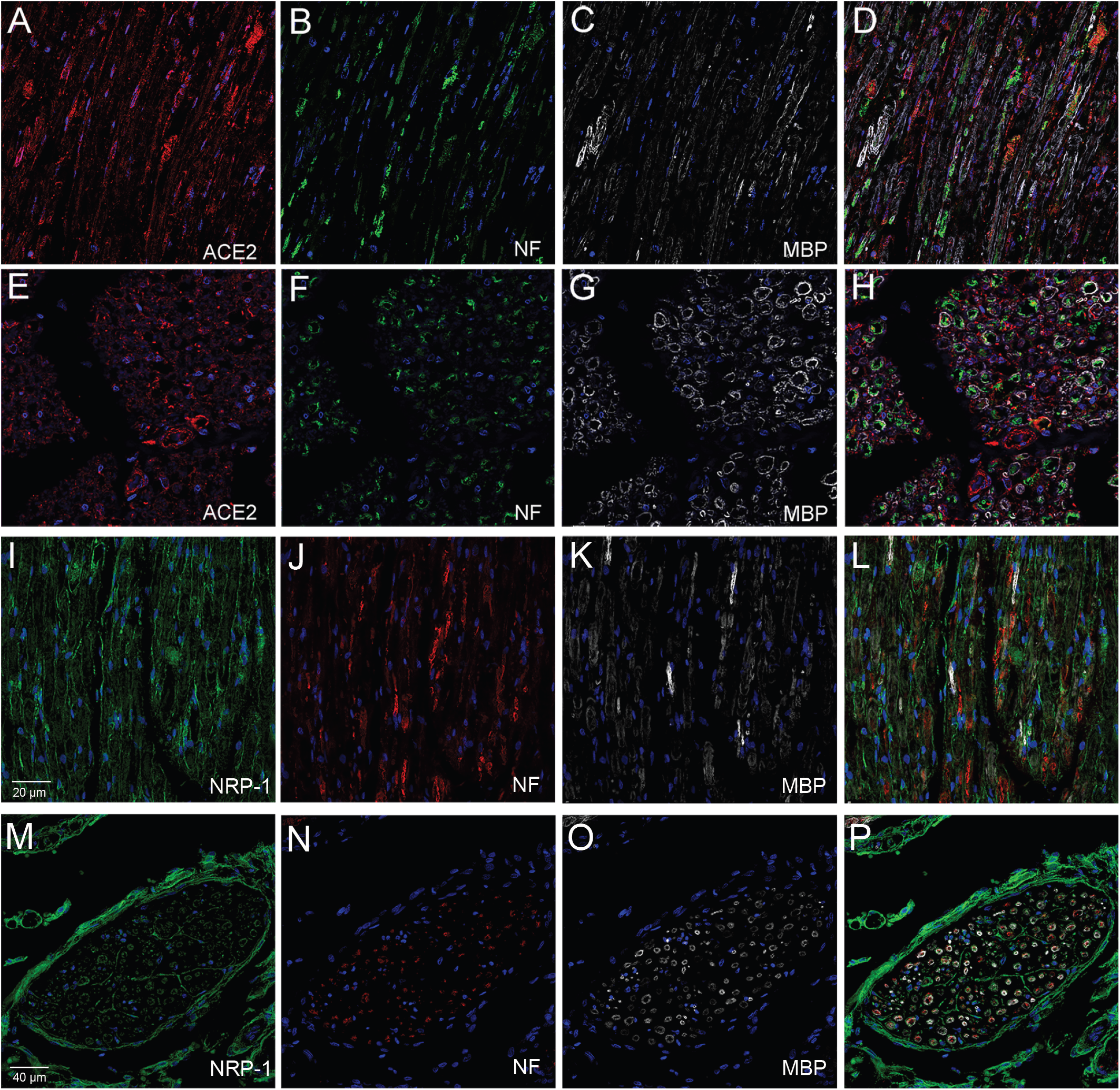
Human glossopharyngeal/vagal nerve at the level of medulla oblongata. The tissue contained both cross sectional cuts of the nerve (E-H & M-P) and longitudinally cut sections of the nerve (A-D & I-L) Six microns thin sections were stained for neurofilament using a chicken primary antibody (Diluted 1:10000) (B, F, J, N) followed by either anti-chicken Alexa-488 (green) (B&F) or anti-chicken Alexa-555 (Red) (J&N). Sections were stained next for ACE2 using a rabbit primary antibody (Diluted 1:100) and an anti-rabbit Alexa-594 (red) conjugated secondary antibody (A&E) or an anti-neuropilin1 (NRP-1) (B) pre-conjugated to Alexa-488 antibody (Diluted 1:100) (I&M). Finally, all sections were stained using an anti-Myelin Basic Protein (MBP) (C, G, K, O) pre-conjugated to Alexa-647 (Diluted 1:100) (colored white). (D, H, L, P) shows the overlay of the three antibodies. Cell nuclei are stained with DAPI (blue). *ACE2 & NRP-1 antibodies were applied overnight, MBP was applied for two hours. *All secondary antibodies were diluted 1:500.

Using small pieces of the frozen human postmortem, we performed PCR to confirm the presence of mRNA of the viral entry sites and the TMPRSS2 protease. We used human lung tissue as a positive control and no template as a negative control. We used two primer pairs for both TMPRSS2 and ACE2 based on literature data [14,15]. All mRNAs were present in both the human lung and the IX/Xth nerve sample (Fig. 5).

### Cultured human cell lines

As described in the Methods section we obtained human cell lines from ATCC to confirm our findings in the postmortem tissues. Both the oligodendrocytes (Fig.4 A-C) and the fibroblasts (Fig. 4D-E) were stained by ACE2, NRP1 and the TMPRSS2 antibodies. ACE2 showed a very fine punctate staining in the membranes and cytoplasm. The NRP1 antiboy stained larger granules in the oligodendrocytes and gave more pronounced membrane staining in fibroblasts.

**Fig.4.**
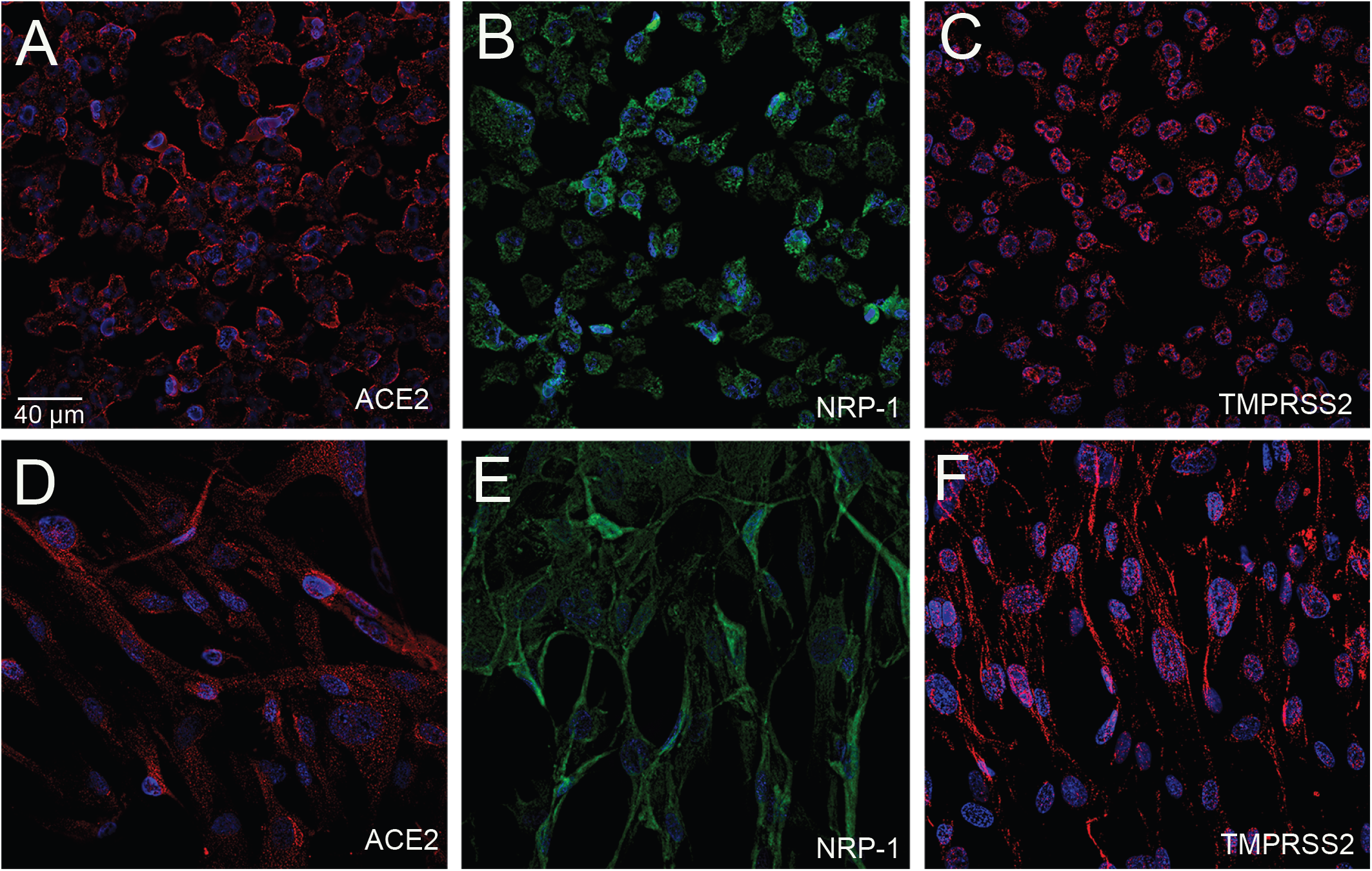
Cultured oligodendrocytes and CNS fibroblasts stained for ACE2, NRP-1 and TMPRSS2 using human immortalized oligodendroglioma cells and human primary brain vascular fibroblast. Chamber slides of each cell line were stained with either ACE2 (A, D) using a rabbit primary antibody and an Alexa-594 (red) conjugated secondary antibody, TMPRSS2 (C, F) using a rabbit primary antibody and an anti-rabbit Alexa-594 (red) conjugated secondary antibody, or an anti-Neuropilin (B, E) pre-conjugated to Alexa-488 antibody. Cell nuclei are stained with DAPI (blue). *All primary antibodies were applied overnight with a 1:100 dilution. *All secondary antibodies were diluted 1:500.

**Fig.5.**
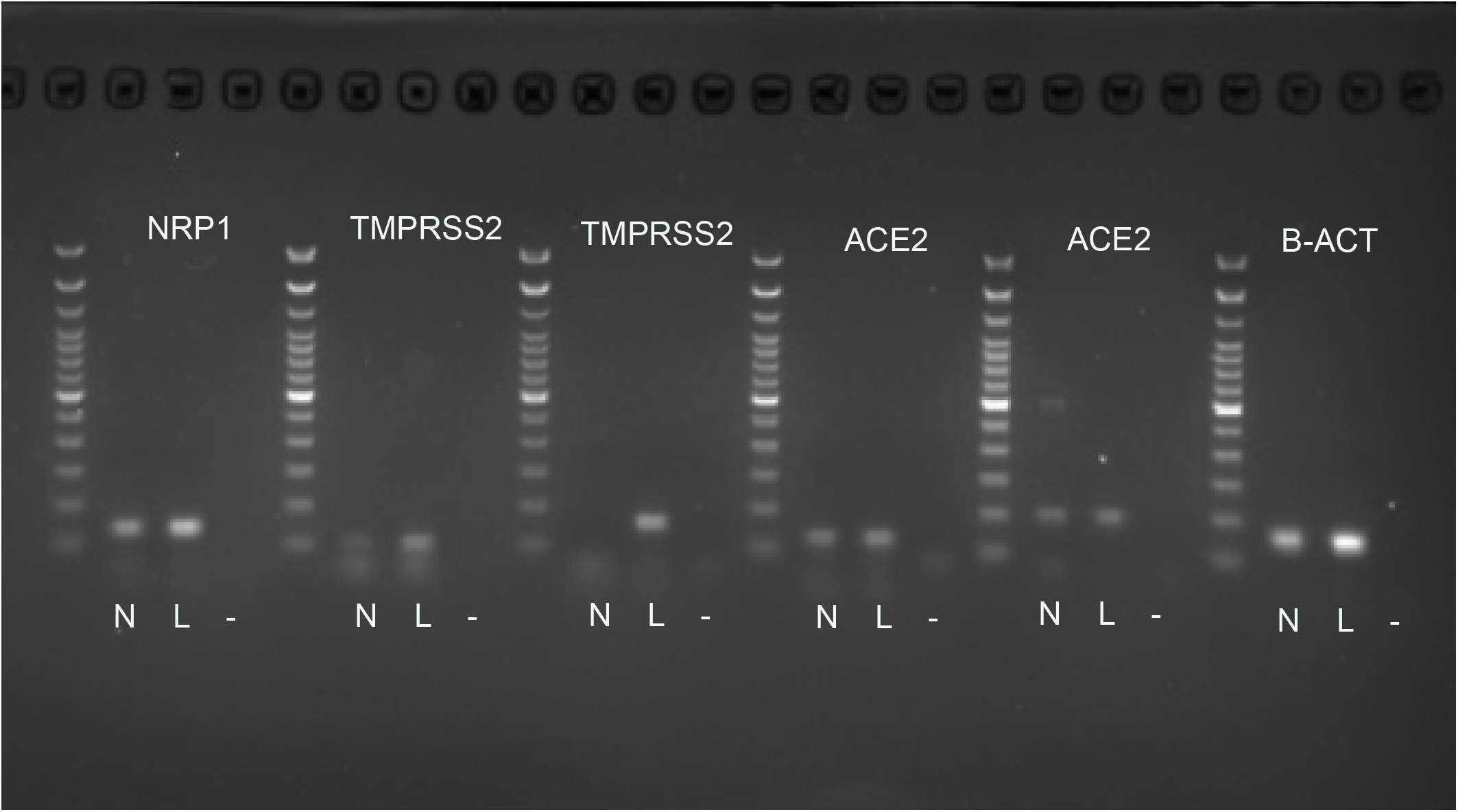
RT-PCR of human frozen nerve. Fresh human nerves (N) were collected and treated with Trizol for RNA preparation. RNA was reverse transcribed for the nerves and human lung (L) RNA purchased as a positive control. RT-PCR was performed to test the presence of NRP1(137bp), TMPRSS2 (106bp and 158bp) and ACE2 (124bp, and 199bp). For normalization, β-actin was used as a housekeeping gene (184bp), and a 100bp ladder was used for size verification. Two primer sets were used for both the ACE2 and the TMPRSS2 [10]. Negative control (-) (no template) does not show any bands.

## DISCUSSION

In the last two years information about the SARS-CoV-2 virus has expanded dramatically. The “neuroinvasive capacity” of the virus was described and in human organoids this invasion was prevented by ACE2 antibodies as well as by CSF isolated from Covid patients [16]. At the start of the pandemic, one of the first widespread observations was that SARS CoV-2 infections result in temporary loss of smell and taste (anosmia and ageusia), even in people who were otherwise asymptomatic. In fact, anosmia and ageusia appear to be the most prevalent manifestations of neuronal disease in COVID-19 with frequencies ranging from 33% to 88% of victims [17]. In addition to loss of taste and smell, inflammatory demyelinating polyneuropathy and Guillain-Barré syndrome (GBS) have also been reported in Covid-19 infections [18]. SARS-CoV-2 seems to have neurotropism and can trigger demyelination [19] by directly injuring myelin-producing cells or causing inflammation.. Neurotropic viruses have long been known to find their way to the brain by means of retrograde transport along axons. The best known and studied among these are the herpes viruses. The ability of these viruses to be taken up by nerve terminals and moved to cell bodies has been exploited in numerous tract tracing studies. Rangon et al. suggested that the SARS-CoV-2 virus invades the vagal nerve and travels from the lung to the brainstem autonomic centers [20]. In addition, genetically manipulated viruses have been used to map neural pathways because they are capable of anterograde and retrograde trans-synaptic migration [21,22]. Other authors have suggested that coronaviruses [23], including the SARS-CoV-2 virus [24,25], enter the CNS via the nasal cavity and olfactory system. Olfactory neurons are uniquely suited to conveying viruses from the periphery to the brain. They are bipolar cells with dendrites that face the external surface of the cribriform plate and axons that traverse the cribriform foramina and terminate in the olfactory bulbs. Once a virus enters neurons in the olfactory bulb it may be able to make its way to cortical areas, including the piriform “primary olfactory” cortex [23]. However, in a recent review Butowt [26] suggested that the evidence for an olfactory route of CNS infection by SARS-CoV-19 is weak and the progression and timing of CNS disease following infection suggest alternative routes, such as spread through vasculature and crossing the blood-brain barrier. Supporting that hypothesis in their studies Lee et al. performed postmortem high resolution magnetic resonance imaging of brains of patients suffering fatal Covid-19 infections and showed widespread microvascular injuries throughout the CNS[27].

In spite of the fact that the loss of taste is just as prominent as the loss of smell in COVID-19 infections, no studies of human tissues have been published that shed light on the mechanisms responsible for ageusia – most likely because of the difficulty of obtaining appropriate cranial nerve samples. One hypothesis is that the loss of taste is secondary to the loss of smell, but most people can distinguish these problems from one another. Furthermore, the gustatory (taste) pathway is very different from the olfactory one [28]. Taste cells are mainly localized in the tongue in papillae surrounded by epithelial cells. The anterior two-thirds of the tongue is innervated by the facial (VIIth) nerve, the posterior third, the throat, the tonsilles and the anterior part of the pharynx are innervated by the glossopharyngeal (IXth) nerve, while the rest of the pharynx and the epiglottis are innervated by the vagal (Xth) nerve. The pseudounipolar cell bodies that give rise to the facial, glossopharyngeal, and vagal nerve fibers are located in the geniculate, petrosal and the nodose ganglia, respectively (Fig.6 B). Their proximal projections terminate in the nucleus of the solitary tract (NTS) in the dorsal medulla (Fig.6 B). All taste signals converge there and incoming signals ascend to the thalamus and eventually reach the primary gustatory center in the insular cortex. The insular cortical neurons project to the medial prefrontal cortex where taste signals are recognized by activation of gustatory working memories, the orbitofrontal cortex where motivational and emotional responses to tastes occur, and ultimately the medial parietal cortex (precuneus) where taste sensations are understood, individually [29]. It should be mentioned that a significant fraction of the fibers in the glossopharyngeal and vagal nerves carry pain, temperature, and pressure signals from the tongue and the upper and lower respiratory areas, including the lungs. All of these sites could also be infected by COVID-19 since the virus can travel from the gustatory and non-gustatory sensory ganglion cells to the medulla to terminate in the spinal trigeminal nucleus. From there fibers ascend to the thalamus and then to the primary sensory cortex (“trigemino-thalamo-cortical pathway”). All of these brainstem and cortical areas mentioned above could potentially be infected trans-synaptically after the virus finds its way into the three taste-receptor containing cranial nerves.

**Fig.6.**
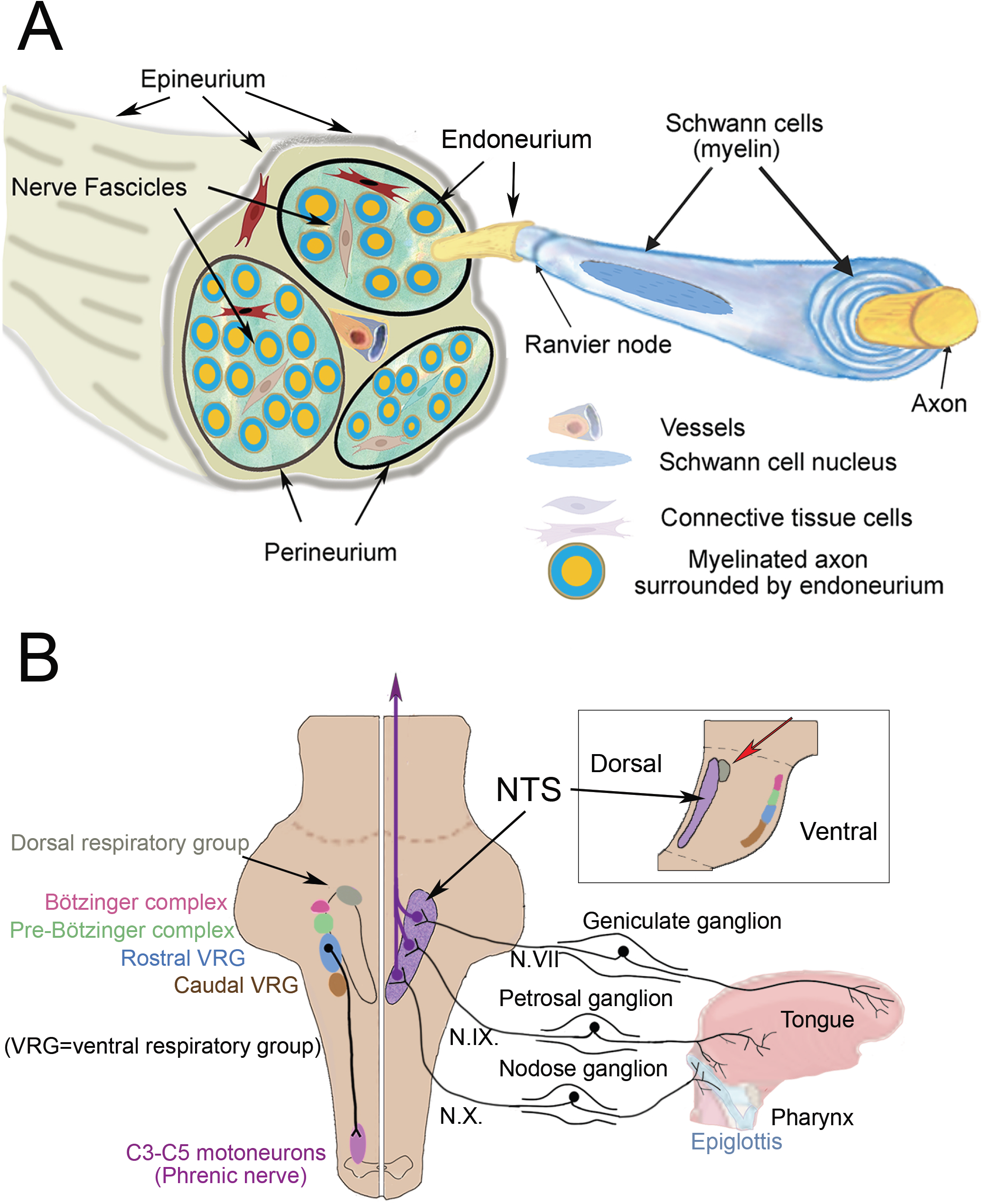
A: Schematic drawing of a human nerve bundle showing possible membranes and spaces where the virus can potentially move along the nerve in a retrograde fashion. B: shows a simplified version of the brainstem from a dorsal view (not indicating the dorso-ventral depth) where the solitary tract neurons receive input from the periphery from the ganglia of the VII, IX and X cranial nerves. The right side shows the intimate closeness of neurons of the pre-Bötzinger and Bötzinger respiratory complex and other respiratory centers, including the cells innervating the phrenic nerve, that could potentially all be harmed by viral invasion through these nerves – including (but not restricted to) the gustatory fibers. The small inset shows a sagittal schematic of the medulla to demonstrate how close the dorsal respiratory group is to the NTS (these two nuclei are labelled with a red arrow in the Figure.

In our study we looked at the vagal and glossopharyngeal nerves in human postmortem samples (non SARS-CoV-2 infected) at the point where the nerves leave (or connect to) the medulla. We found that both of the binding sites for the SARS-CoV-2 viral spike proteins, ACE2 and NRP1, are widely expressed in the nerve bundles, in myelin sheaths as well as axons. Vascular walls also contain these entry sites. Cranial nerves contain a large number of connective tissue cells (Fig.6 A) that form sheaths that surround the nerve (epineurium), the fascicles (perineurium), and the myelinated axons (endoneurium) [30,31]. The function of these fibroblast-like cells is not clear. Many of the ones in human cranial nerves are also immunopositive for lymphatic endothelial markers and are in contact with the endoneurial fluid suggesting a role in immune survaillance [6]. Unlike peripheral fibroblasts (which are mesodermal in origin) these cells are derived from the neural crest. Some of them make nestin and may be involved in regenerating nerves and myelin following injuries [31,32]. We found both SARS-CoV-2 viral entry sites in all cell types within the nerve. We also found TMPRSS2, the protease that primes the spike protein within the infected cell. TMPRSS2 has been shown to facilitate infections that can be blocked by protease inhibitors [12]. TMPRSS2 expression was high in fibroblasts, but is present in all cell types within the nerve. Vascular endothelium and smooth muscle cells expressed lower levels of TMPRSS2 than they did ACE2.

Given these results, we tried to imagine how all these cellular elements might play a role in the loss of taste in infected individuals. The gustatory pathway originates in the oral cavity where ACE2 has been shown to be present in human taste buds [33] and the tongue epithelium [34]. After it infects taste-sensing nerve endings the virus might make its way to the CNS where there are various ways for neurotropic viruses to infect neural tissues see [10]. The SARS-CoV-2 virus could bind to papillae in the tongue; enter the neurons (since there is no blood brain barrier in the periphery); and move along the axon, highjacking the transport routes used physiologically by the neuron to carry particles as other viruses do [22,24,35]. Alternatively, a “trojan horse” mechanism [36] could be involved: the virus could “hitch a ride” in cells like lymphocytes or macrophages and travel along the axons to the CNS. Given the presence of T cells and macrophages in the human trigeminal nerve and ganglion this seems feasible. Lymphatic endothelial marker positive fibroblasts might also participate in carrying the virus within fluid compartments of nerve bundles (endoneurial fluid) using the arterial pulsation as a driving force [4]. The presence of all the components necessary for infection in neuronal connective tissue raises the possibility that the virus might infect fibroblasts (the nerve sheaths) and later have a “transient adhesive interaction” with lymphocytes to spread the virus, as mentioned in an earlier review [35]. Last but not least, the virus could enter from the circulation and infect any or all elements of the nerve. Once the virus reaches the solitary tract complex in the medulla, the area where the sensory neurons of the IXth and Xth nerves are found, it is in very close proximity to neurons that regulate respiratory functions in humans (Fig.6 B). The glossopharyngeal and vagal nerves may transfer the virus to medullary respiratory nuclei including the dorsal respiratory nucleus (the only dorsal respiratory cellgroup close to the nucleus of the solitary tract (NTS)), the pre-Bötzinger and Bötzinger complexes, and two additional ventral nuclei located in the caudal ventrolateral medulla (Fig.6 B). The only dorsal respiratory group in the medulla is in the immediate vicinity of the NTS (labelled in Fig.6B is thought to be involved in inspiratory function [37]. The pre-Bötzinger complex acts as a respiratory pacemaker regulating the rhythm of breathing [37-40]. The two ventral respiratory cell groups project to to cervical segments 3-5 where the phrenic nerve fibers that innervate the diaphragm arise. Damaging these cells could have a significant effect on survival making it very difficult for patients to breathe on their own after they have recovered from pneumonia and assisted breathing. In fact, a COVID19 patient has been described who had severe widespread damage to his neurons, axons, glia and myelin sheath. Electron microscopy revealed particles “referable to virions of SARS-Cov-2” [41]. The case report states that “Upon withdrawal of sedation and paralysis the patient became profoundly agitated with severe ventilator asynchronies such as ineffective efforts and double cycling, that required deep sedation, and paralysis. The patient died after 19 days.” [41]. In another paper analyzing neuropathology of patients who died following infection with SARS-Cov-2 one patient is mentioned where the virus seemed to be present in individual cells of cranial nerves [42]. A review was recently published summarizing data on the role of neuro-invasion of brainstem respiratory centers by the SARS-Cov-2 in the respiratory failure of some patients [43].

We earlier talked about viral proliferation in the olfactory epithelium and subsequent invasion of the cells that ensheath olfactory neurons. The ACE2 entry site was also shown to be present in non-neuronal cells of the olfactory epithelium [25], just as we found ACE2, NRP1, and TMPRSS2 in the supportive cells of glossopharyngeal and vagal nerves. In this regard, it is interesting that 40 years ago olfactory bulbectomy in rats was reported to produce depression-like symptoms in the animals [44]. COVID-19 infections are commonly associated with depression in people [45]. The two findings could be related and it would be interesting to know whether there is a correlation between loss of smell and depression in Covid patients.

It should be clear that there is still a lot to learn about how the SARS-Cov-2 virus affects the peripheral and central nervous system. Demonstration of the expression of molecules that it can use as an entry site is just the beginning. We need to find out how it choses cells to infect, how it highjacks neuronal transport mechanisms (anterograde and retrograde), how it uses non-neuronal (fibroblast and immune) cells, and how it evades immune surveillance within the nerve and CNS. We know that eventually a healthy immune system eliminates the virus, because taste usually returns within a matter of weeks, but it will be important to learn more about viral entry to stop CNS infections completely.

## Acknowledgement

This study was supported by the Intramural Research Program of the National Institute of Dental and Craniofacial Research (NIDCR), NIH (intramural project no. ZDE000755-01), and the Human Brain Tissue Bank, Semmelweis University, Budapest, Hungary. In Hungary, the brains of donors were dissected and brainstem with cranial nerves were isolated. The tissues were either flash frozen or embedded in paraffin after formalin fixation. M.P. is supported by grant from the Hungarian Brain Research Program (2017-1.2.1-NKP-2017-00002). The study was approved by European ethics regulations under permission TUKEB #189/2015.

The authors thank Magda Toronyay-Kasztner for her assist with preparing the manuscript and Dr. Michael Brownstein for expert editing work.

## Notes

### Competing Interest Statement

The authors have declared no competing interest.

### Summary of Updates

Added more recent literature to discussion; expanded Fig.6

